# Colonic epithelial regeneration shapes susceptibility to *Clostridioides difficile* infection

**DOI:** 10.64898/2026.05.21.727036

**Authors:** Adrianne D. Gladden, Paola Zucchi, Albert Tai, Rebecca Batorsky, Carol A. Kumamoto

**Affiliations:** Department of Molecular Biology and Microbiology, Tufts University, Boston, Massachusetts, 02111, USA; Graduate School of Biomedical Sciences, Tufts University, Boston, Massachusetts, 02111, USA; Department of Immunology and TUCF Genomics Core, Tufts University, Boston, Massachusetts, 02111, USA; Data Intensive Studies Center, Tufts University, Medford, MA, 02155

## Abstract

*Clostridioides difficile* infection (CDI) susceptibility and severity are strongly associated with preexisting colonic inflammation. However, chronic inflammatory conditions such as cystic fibrosis rarely progress to symptomatic CDI despite high rates of *C. difficile* colonization, suggesting that inflammation alone is insufficient to explain disease vulnerability. Notably, populations relatively protected from symptomatic CDI exhibit impaired regenerative capacity within the colon epithelium. Here, we used single cell RNA sequencing of human colonoid monolayers to map markers of CDI susceptibility and severity to cell populations associated with inflammation and epithelial repair. We identified an inducible microfold-like (M-like) population that is largely absent from the healthy colon but emerges during inflammation and regeneration. These cells were enriched for markers of severe CDI, *C. difficile* toxin interaction genes, and elevated CCL20 and CFTR expression. Spatial imaging localized CCL20-producing cells to wound-like gaps in mock and CDI-treated colonoids, identifying a repair-associated niche active independent of infection. Following exposure to *C. difficile*, wound-healing transcription within the M-like lineage declined while tuft-like populations expanded and upregulated genes associated with immune cell recruitment. These findings demonstrate that epithelial regeneration shapes host CDI vulnerability.

**IMPORTANCE:** *Clostridioides difficile* infection can lead to severe illness and death in vulnerable populations despite available treatments. Clinical signs of inflammation during active *Clostridioides difficile* infection are strongly associated with disease outcome, yet these responses primarily reflect tissue damage already underway, limiting opportunities to prevent progression. In contrast, conditions linked to severe disease, including inflammatory bowel disease and antibiotic exposure, are associated with colonic inflammation before infection or at the time of diagnosis, highlighting an opportunity for earlier identification of high-risk individuals. Using human colonoid single cell transcriptomics and spatial imaging, we identified a microfold-like cell population enriched for inflammatory mediators and *Clostridioides difficile* toxin interaction genes linked to severe disease. This population was active even in the absence of infection, suggesting that repair-associated populations within the inflamed colon may help identify susceptibility to severe CDI before clinical progression occurs.

## INTRODUCTION

*Clostridioides difficile* infection (CDI) afflicts millions worldwide each year with a range of symptoms that include diarrhea and varying degrees of colonic inflammation. Clinical disease progression leads to serious illness in 17%, recurrence in over 25%, and death in 6% to 37% of CDI cases in vulnerable populations, despite available treatments.^1–4^ Given that delays in appropriate intervention increase the risk of complications and recurrence, early prediction of disease susceptibility and severity is crucial for improving patient outcomes.^5^

CDI initiates in the colon when a susceptible host is exposed to toxigenic bacteria or when a *C. difficile* carrier becomes susceptible to infection, with symptoms typically presenting within 24-48 hours. Disease onset coincides with a metabolic shift in *C. difficile* promoted by the inflamed colonic environment that enables glycosylating toxin production and amplifies host pro-inflammatory immune responses at damaged sites within the colonic epithelial barrier.^4,6–11^ Consistent with this, previous in vitro and in vivo analyses of colon epithelia during early CDI identified increased expression of genes associated with *C. difficile* toxin interaction, metabolic stress, immune signaling, cell cycle activation, proliferation, and epithelial inflammation.^12–14^ Additional studies demonstrated epithelial barrier disruption and damage to colonic stem cells associated with reduced regenerative capacity.^15,16^ Importantly, CDI severity has been closely linked to the location of toxin interaction within the epithelium: toxin exposure restricted to the epithelial surface resulted in mild disease, whereas severe CDI and poor outcomes occurred when toxins reached the protected stem cell compartment.^16^

Beyond inflammatory responses induced during active infection, elevated colonic inflammation preceding CDI or present at the time of diagnosis is also strongly associated with severe disease and treatment failure. Such inflammatory states are frequently associated with conditions including inflammatory bowel disease (IBD), colon cancer, or other medical issues, along with advanced age and certain antibiotic or other drug exposures.^17–27^ However, progression to clinical disease varies markedly across populations.^25,28–30^ For example, neonates are less likely to experience symptomatic CDI, even with similar antibiotic usage and high rates of *C. difficile* colonization (>70%).^31^ Likewise, individuals with cystic fibrosis (CF) appear to be less prone to clinical infection, despite having increased *C. difficile* carriage (>50%), antibiotic use, and existing colonic inflammation.^32–34^ Notably, both neonates and individuals with CF exhibit impaired regenerative capacity within the colon epithelium. These paradoxes suggest that inflammation alone is insufficient to explain either the occurrence or the severity of CDI and highlight the need to define host determinants that shape disease outcomes.^35,36^

Although dysbiosis has been widely implicated in CDI risk and pathogenesis, high-risk clinical states are also characterized by epithelial injury and inflammation.^37–39^ In IBD, epithelial mitochondrial dysfunction precedes microbiota disruption and promotes inflammatory signaling and tissue damage.^40,41^ The antibiotic clindamycin, among the strongest clinical risk factors for severe CDI and mortality, similarly induces mitochondrial damage in intestinal epithelial cells in a microbiota-independent manner and drives an IBD-like inflammatory state in the colon.^42–46^ Drug-induced mitochondrial injury also sensitizes epithelial cells to *C. difficile* toxin and can increase the likelihood of severe disease.^27^ Thus, epithelial cell populations arising during damage and repair may contribute to CDI vulnerability.

Continuous regeneration of the colonic protective barrier is driven by differentiation of stem cells which reside at the base of crypts. However, when the stem cell niche is damaged, the inflammatory response initiates a multi-step repair process: reversion of mature epithelial cells to a stem-like state (dedifferentiation), cell proliferation, and redifferentiation into specialized cells.^47–50^ Regeneration in this context is accompanied by extensive transcriptional reprogramming and the emergence of epithelial states that are mostly absent from the healthy colon.^51–59^ Such states are characteristic of colon epithelia associated with high risk for severe CDI.^60–62^ The colon exhibits a distinct vulnerability to epithelial damage compared to other intestinal regions, leading to more severe inflammation and barrier dysfunction.^63^ These features may help explain why CDI is largely restricted to the inflamed colon.

To characterize epithelial transcription in response to early CDI, independent of microbiota and immune recruitment, we previously established a germ free human colonoid monolayer model. Using this system, we identified *CCL20* as a prominent toxin-dependent epithelial response to acute CDI.^64^ In this study, we used single-cell transcriptomics across multiple timepoints to determine how markers of CDI susceptibility and severity map to epithelial cell populations associated with regeneration early in infection.

We identified a microfold-like cell population enriched for inflammatory signatures and host factors involved in *C. difficile* toxin interaction. Trajectory analysis resolved upstream low-transcript epithelial populations linked to the emergence of this repair-associated lineage. In response to CDI, progression through this regenerative path was disrupted, resulting in unresolved epithelial damage and a compensatory expansion of the tuft-like cell lineage. Our findings implicate epithelial regeneration in CDI susceptibility and disease progression.

## RESULTS

### Human colonoids capture clinical CDI severity markers within a conserved transcriptional landscape

We used antibiotic-treated human colonoid monolayers that recapitulate key features of the mature colonic epithelium to examine early cellular responses to *C. difficile* infection.^64^ To map signatures of CDI vulnerability and severity to epithelial subtypes, we performed single-cell RNA sequencing (scRNA-seq) on monolayers collected 4, 15, and 25 hours following mock-treatment (“Mock”) or *C. difficile* inoculation (“*C. difficile*-infected”) (Figure 1A).

**Figure 1.**
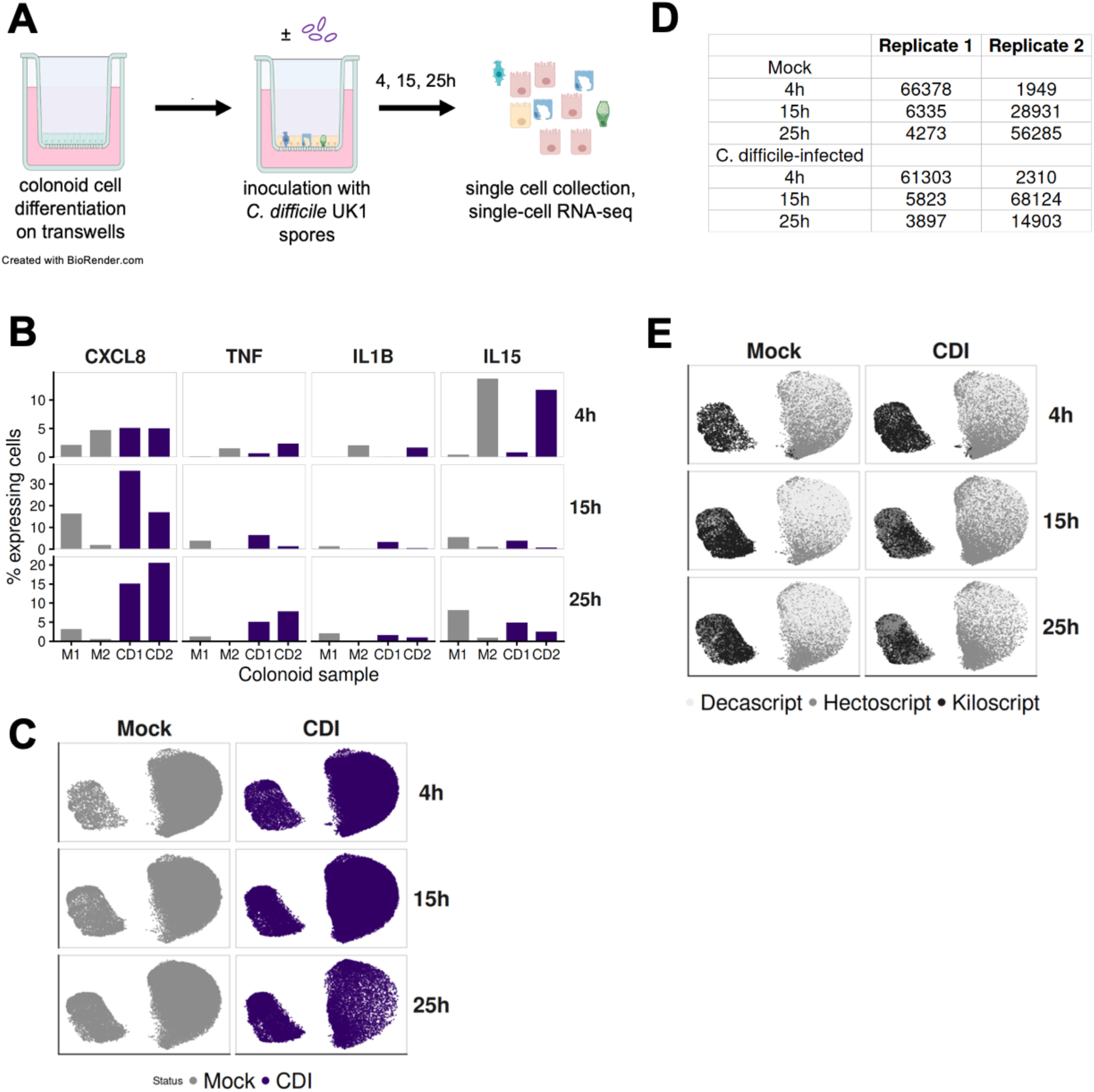
Human colonoids capture clinical CDI severity markers within a conserved transcriptional landscape. (A) Experimental design. Human colonoids were differentiated on Transwell inserts for 5 days, inoculated with *Clostridioides difficile* UK1 spores or mock-treated, and collected for single-cell RNA sequencing at 4, 15, and 25 hours after inoculation. (B) Percentage of cells expressing representative inflammatory genes associated with CDI severity (*CXCL8*, *TNF*, *IL1B*, and *IL15*) in each replicate sample. (C) UMAP visualization of integrated colonoid epithelial cells colored by CDI status. UMAP projection stratified by CDI status and timepoint at 4, 15, 25 h post inoculation. (D) Single-cell profile counts from each biological replicate and timepoint after filtering. (E) UMAP visualization of integrated colonoid epithelial cells colored by transcript-count classes (Decascript, Hectoscript, and Kiloscript). UMAP projection stratified by CDI status and timepoint at 4, 15, 25 h post inoculation.

**Figure S1.**
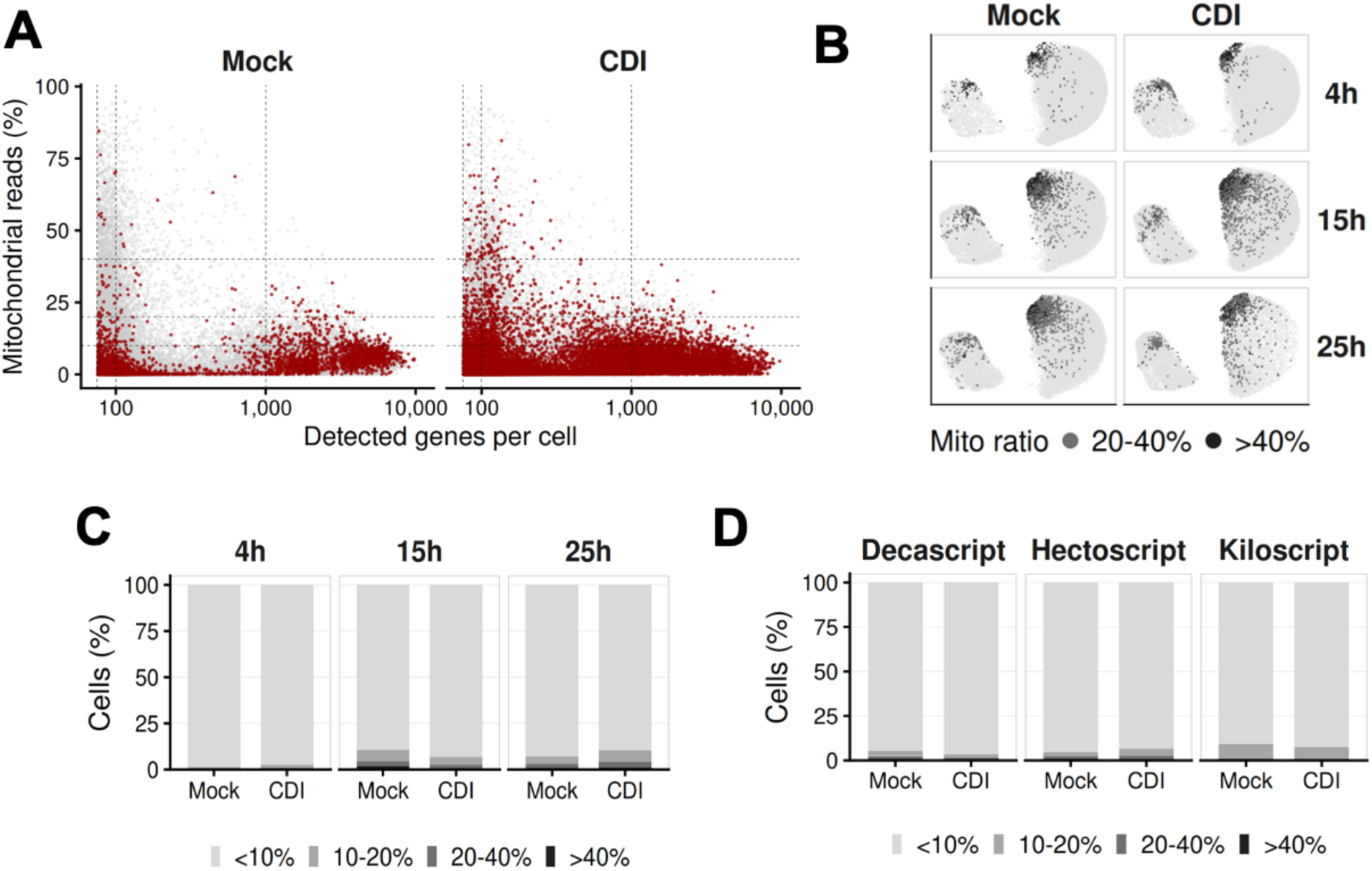
Characterization of single-cell profiles across mitochondrial read fraction and transcript complexity, related to Figure 1. (A) Distribution of detected genes per cell profile and mitochondrial read fraction in mock and CDI colonoid samples. Cells with high (top 10%) CDI-associated inflammatory (*CXCL8*, *TNF*, *IL1B*, and *IL15*) module scores are colored red. Horizontal dashed lines indicate commonly used mitochondrial read fraction filtering thresholds. (B) UMAP visualization of integrated colonoid epithelial cells colored by mitochondrial read fractions (20–40% and >40%). UMAP projection stratified by CDI status and timepoint at 4, 15, 25 h post inoculation. (C) Distribution of mitochondrial read fractions across timepoints and CDI status. (D) Distribution of mitochondrial read fractions across transcript-count classes (Decascript, Hectoscript, and Kiloscript) in Mock and CDI samples.

The clinical relevance of our colonoid model was validated by evaluating expression of established CDI severity markers, including *CXCL8*, *TNF*, *IL1B*, and *IL15*. These transcripts were detectable in both infected and Mock samples, with variation across replicates and timepoints (Figure 1B). Expression of this signature aligns with clinical observations linking colonic inflammation, even in the absence of infection, to CDI susceptibility and outcomes. This biological feature was used to inform subsequent analyses of the colonoid dataset.

We combined single-cell transcriptomes from mock-treated and infected monolayers based on similarity of their gene expression patterns to enable direct comparison between conditions. Mock and infected cells showed substantial transcriptional overlap, with no evidence of distinct infection-specific groupings (Figure 1C). The final dataset comprised 211,537 single-cell transcriptomes spanning a broad range of detected gene counts (Figure 1D). Because reduced transcript complexity has previously been linked to regenerative dedifferentiation, low-transcript cells were retained for downstream analysis.^65^ We applied the previously described Hectoscript (100–999 genes) and Kiloscript (≥1000 genes) classifications and extended this framework to include a lower-transcript category, Decascript (76–99 genes).^65^ UMAP visualization of these transcript-count revealed a continuum of transcriptional states rather than discrete populations (Figure 1E).

Finally, Because colonic inflammation has been associated with altered mitochondrial gene expression, we evaluated the relationship between inflammatory signaling and mitochondrial read proportion.^13,40,66,67^ Cells expressing CDI-associated inflammatory signatures were distributed across a broad range of detected gene counts and mitochondrial read proportions in both mock and infected, similar to the overall cell population (Figure S1A). A subset of cells with high mitochondrial content (>40%) localized to a distinct transcriptional cluster (Figure S1B), while most cell profiles had <10% mitochondrial reads. These distributions remained consistent across transcript-count classes and across all infection timepoints (Figure S1C-D).

### Reference-based cell type labels reveal inflamed regenerative epithelial states with and without CDI

We subsequently established a strategy to assign epithelial cell types to the colonoid transcriptomes in the context of inflammatory activation (Figure S2A). SingleR,^68^ an automated program for reference-based and cluster-independent annotation, was used with a combined reference composed of three publicly available single-cell atlases from the same study.^56^ These atlases included epithelial transcriptomes from colon biopsy samples of healthy control subjects (n = 12), and paired samples from inflamed regions (“Inflamed IBD Colon”) and non-inflamed regions (“Non-inflamed IBD Colon”) of subjects with inflammatory bowel disease (n = 18). Most colonoid cells correlated strongly with the inflamed IBD reference regardless of CDI status or timepoint (Figure 2A). To evaluate this annotation strategy, we re-analyzed published epithelial single-cell datasets from healthy and IBD colon^58^ and antibiotic-treated colonoids.^20^ Datasets associated with CDI risk factors, including inflamed IBD epithelia and antibiotic-treated colonoids, showed the highest correlation with inflamed IBD epithelial reference subtypes (Figure S2B). Based on these results, we made final cell type predictions for this study dataset using SingleR with the inflamed IBD colon reference only.

**Figure 2.**
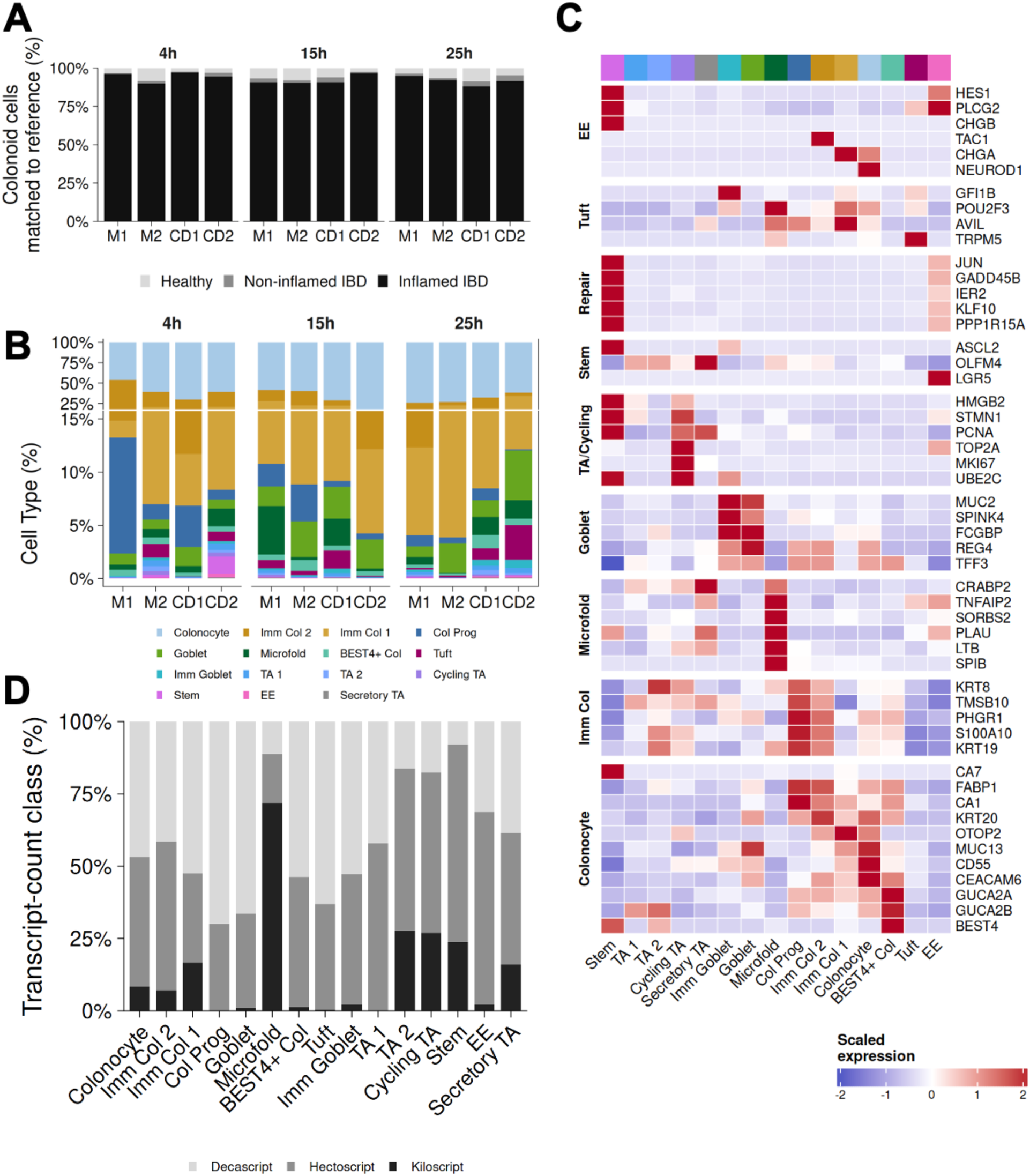
Reference-based cell type labels reveal inflamed regenerative epithelial states with and without CDI. A) Percentages of colonoid cells by timepoint and replicate annotated with reference type in combined atlas of epithelial single-cell profiles from healthy, non-inflamed IBD, and inflamed IBD human colon tissue. (B) Epithelial cell-type composition in each sample at all timepoints. Stacked bars show the proportion of cell types within each sample. A broken y-axis is used to visualize lower-abundance populations while preserving the dominant colonocyte population. (C) Curated marker heatmap supporting epithelial state annotation. Heatmap shows scaled average expression of representative genes across annotated cell types. (D) Distribution of transcript-count classes (Decascript, Hectoscript, and Kiloscript) across cell types.

**Figure S2.**
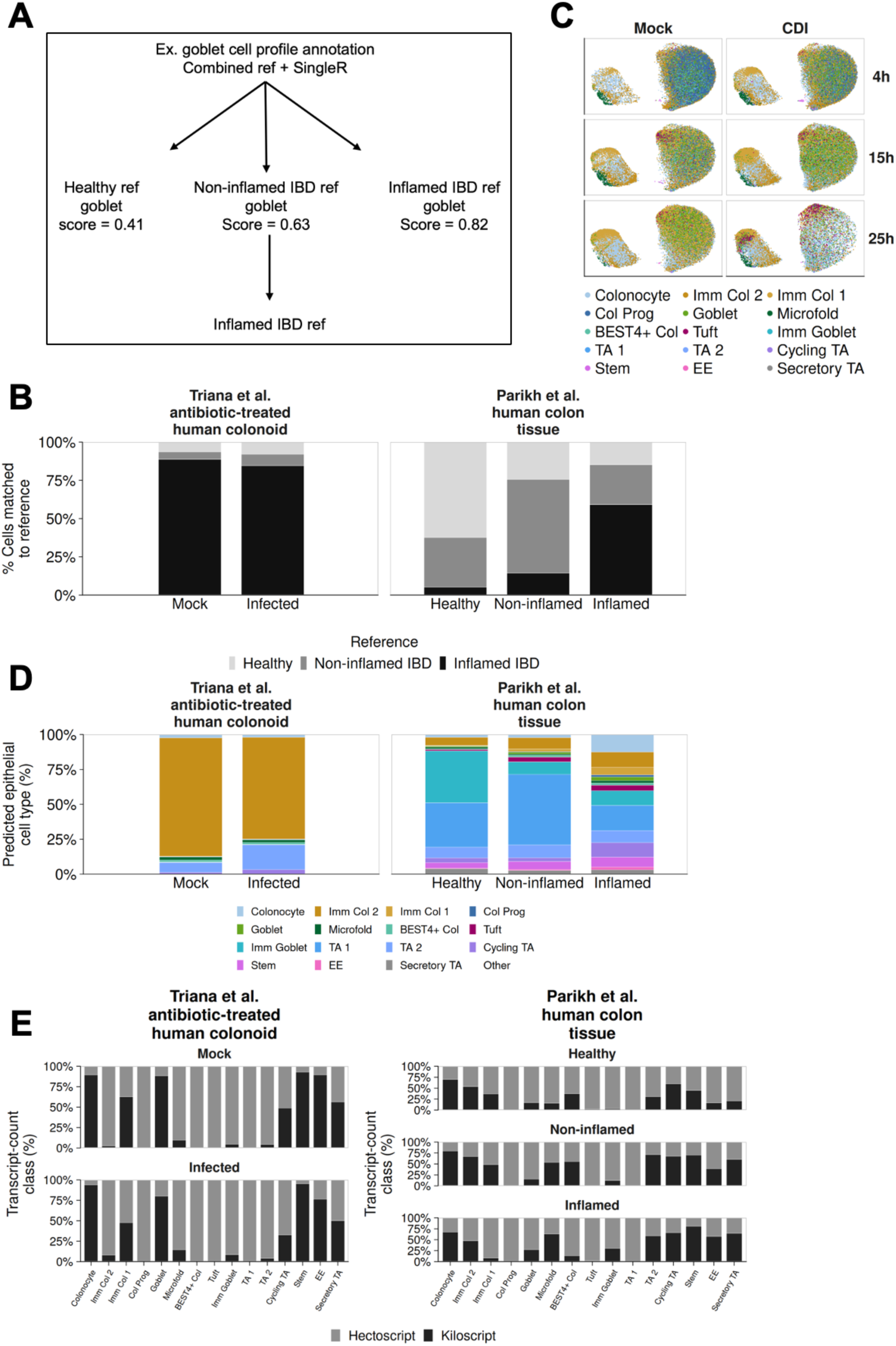
Disease-contextual epithelial type annotation and external dataset validation, related to Figure 2. (A) Schematic of the cell type annotation workflow used to identify reference type for final predictions. SingleR annotation was first performed using a combined atlas of epithelial single-cell profiles from healthy, non-inflamed IBD, and inflamed IBD human colon tissue from Smillie et al. Example similarity scores for a query goblet cell profile are shown schematically. The dominant disease-state reference is identified for each dataset and then used for downstream epithelial cell type annotation. (B) Reanalyzed external datasets used to validate method for reference based cell type annotation. Triana et al. antibiotic-treated human colonoids and Parikh et al. human colon tissue samples were annotated using Singler with the combined reference. Bars represent proportions of cells annotated by each disease-state reference for each sample. (C) UMAP visualization of all samples in this study integrated. Points representing single-cell profiles are colored by cell types annotated using the inflamed IBD colon reference. UMAP projection stratified by CDI status and timepoint at 4, 15, 25 h post inoculation. (D) SingleR-predicted cell-type composition in external validation datasets following disease-state reference cell type annotation. Bars represent the proportion of cells assigned to each epithelial subtype within each sample. (E) Distribution of transcript-count classes across predicted epithelial cell types in external validation datasets. Bars represent the proportion of Hectoscript and Kiloscript cells within annotated epithelial subtypes for each sample.

We identified 15 epithelial subtypes (Figure 2B and Figure S2C), including mature, immature, and progenitor colonocytes, together with goblet, stem, and transit-amplifying (TA) populations. We also detected cell types that are rare in the healthy colon but characteristic of the inflamed colon, including BEST4+, tuft-like, enteroendocrine (EE), and microfold-like (M-like) cells.

Expression of curated canonical and inflammation-associated markers supported the predicted epithelial identities (Figure 2C). Most prevalent were colonocytes expressing lineage markers (*KRT20*, *FABP1*) in parallel with a conserved program of inflammatory damage markers, including *MUC13*, *CD55*, and *CEACAM6*. Within the progenitor subset, we observed a stem population expressing canonical markers (*OLFM4*, *ASCL2*) alongside a robust injury-response signature (*GADD45B*, *JUN*) and the epithelial-to-mesenchymal transition (EMT) mediator *HES1*. This profile is characteristic of damage-associated regenerative stem cells (DARSCs), which emerge during epithelial repair. The secretory lineage comprised goblet cells (*MUC2*, *TFF3*) and M-like and Tuft-like populations that shared the inflammatory-sensing marker *TNFAIP2* in addition to their canonical markers (*SPIB* and *POU2F3/TRPM5*, respectively). M-like cells also co-expressed immature colonocyte markers, as expected for inducible M-like cells. We also observed *LGR5+* enteroendocrine cells (Figure 2C) previously associated with reserve stem-like activity during epithelial wound repair and inflammation. EE cells were most abundant at the 25h CDI timepoint (Figure 2B). Epithelial subtype distributions in our dataset were similar to those observed in external colonoid and human colon tissue datasets, including increased immature colonocyte and tuft cell populations within inflamed IBD-associated epithelia (Figure S2D).

Transcript-count classes differed substantially across predicted epithelial populations, with tuft and colonocyte progenitor populations enriched for low-gene Decascript and Hectoscript cells, whereas stem-like and M-like populations were enriched for Kiloscript cells (Figure 2D). Similar transcript-count distributions were observed across external colonoid and human tissue datasets, supporting reproducible associations between transcript complexity, epithelial identity, and inflammatory context (Figure S2E).

### Trajectory analysis reveals a damage-associated regenerative path and a distinct epithelial lineage

Cells were grouped into clusters which were named based on cell type composition together with transcript complexity. We defined 8 clusters including regenerative progenitor (stem/TA), M-like, secretory, colonocyte I/II/III, mature colonocyte, and tuft-like (Figure 3A). Mock samples were used to infer baseline epithelial progression without CDI-associated disruption of these trajectories. Two epithelial lineages emerged from low-transcript colonocyte populations: an M-like lineage and a tuft-like lineage (Figure 3B-C). The M-like lineage progressed through regenerative progenitor and colonocyte clusters before reaching the M-like state, whereas the tuft-like lineage showed more limited progression and remained enriched for lower-transcript colonocyte and tuft-like populations (Figure 3B-C). RNA velocity broadly supported the inferred direction of these trajectories (Figure S3B).

**Figure 3.**
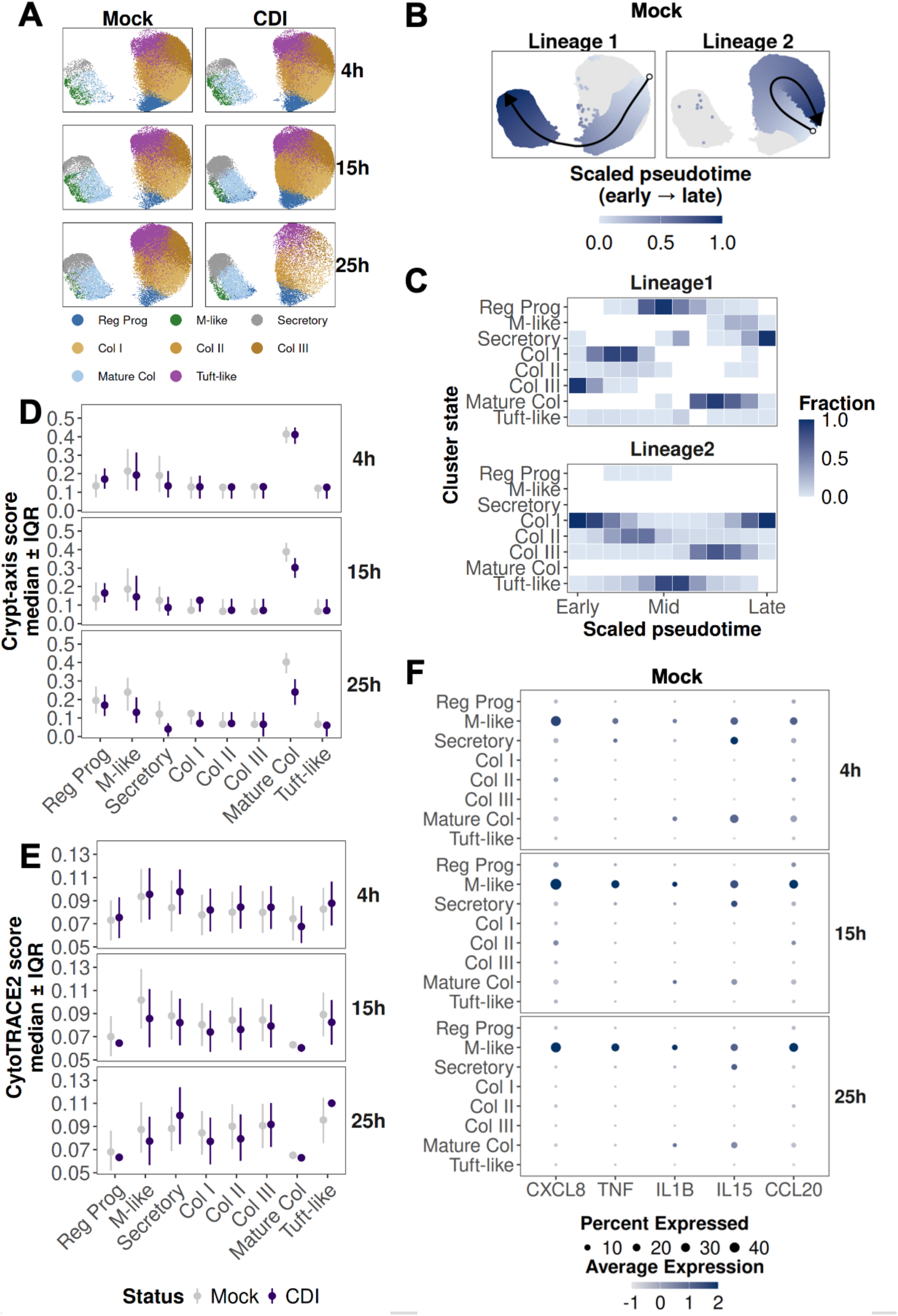
Trajectory analysis reveals a damage-associated regenerative path and a distinct epithelial lineage. (A) UMAP of integrated colonoid epithelial cells colored by cell state cluster and shown separately by condition (Mock, CDI) and timepoint (4 h, 15 h, 25 h). Cells are grouped into regenerative progenitor, M-like, secretory, colonocyte-biased (Col I–III), mature colonocyte, and tuft-like states. (B) Lineage inference in Mock samples projected onto the UMAP manifold. Two lineage trajectories are shown with points colored by scaled pseudotime (early to late). Arrows indicate inferred directionality along each lineage. (C) Distribution of cell states in Mock samples across pseudotime for each lineage. Each tile represents the fraction of cells assigned to a given state within binned pseudotime intervals. (D) Crypt-axis scores and (E) CytoTRACE2 scores for each cell state stratified by timepoint and CDI status. Points indicate medians and error bars indicate interquartile ranges for each cell state. (F) Dot plot of *CXCL8*, *TNF*, *IL1B*, *IL15*, and *CCL20* expression across cell states in Mock samples at each timepoint. Dot size indicates the percentage of cells expressing each gene and color indicates scaled average expression.

**Figure S3.**
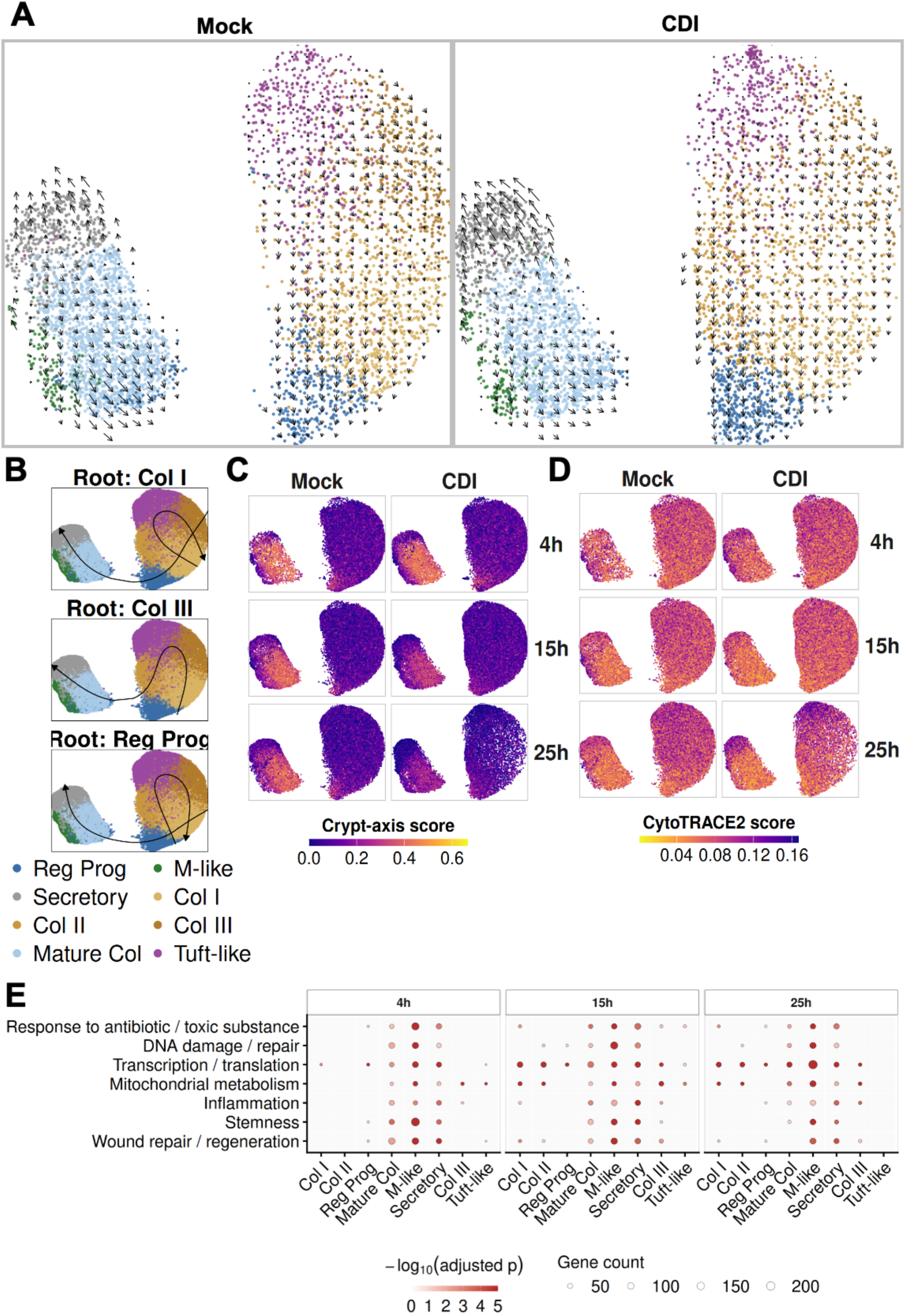
Crypt-axis gradients, CytoTRACE2 scores, lineage structure, and RNA velocity across Mock and CDI colonoids, related to Figure 3. (A) RNA velocity streamlines overlaid on UMAP embeddings for Mock and CDI colonoids at each timepoint (4 h, 15 h, 25 h). Arrows indicate the direction and magnitude of inferred transcriptional change. Cells are colored by cell state. (B) Lineage inference in Mock samples using alternative root clusters (Col I, Col III, and regenerative progenitor). Inferred trajectories are projected onto the UMAP manifold with arrows indicating inferred directionality. (C) UMAP visualization of integrated colonoid epithelial cells colored by crypt-axis score and shown separately by condition (Mock, CDI) and timepoint (4 h, 15 h, 25 h). (D) UMAP visualization of integrated colonoid epithelial cells colored by CytoTRACE2 score and shown separately by condition (Mock, CDI) and timepoint (4 h, 15 h, 25 h). (E) Representative Gene Ontology (GO) categories enriched among genes identified in mock-treated epithelial states across timepoints that were differentially expressed in the same direction in both replicates. Dot size indicates the number of overlapping genes and color intensity represents −log10 adjusted p value.

Crypt-axis and CytoTRACE2 analyses were used to independently estimate differentiation state and developmental potential, respectively (Figure 3D-E and Figure S3C-D). Mature colonocytes were predicted to be the most differentiated overall, although this cluster showed intermediate crypt-axis scores and directly preceded the M-like state along the inferred trajectory. This organization aligns with prior observations that inflamed or damage-associated colonic M-like cells are induced through colonocyte transdifferentiation during epithelial repair. Developmental potential peaked in M-like cells under Mock conditions and in tuft-like cells during CDI despite major differences in transcript complexity between these populations.

To further characterize transcriptional programs associated with these lineages without infection, we performed marker analysis and gene ontology (GO) enrichment using mock colonoids only (Figure S3E; Tables S1A-G and S2). The M-like lineage, including regenerative progenitor and transitional states, was enriched for programs related to antibiotic and toxic substance response, DNA damage and repair, transcriptional/translational changes, mitochondrial metabolism, inflammation, stemness/plasticity, and wound repair. By contrast, the tuft-like lineage showed more limited enrichment, primarily involving transcriptional/translational and mitochondrial metabolic programs with minimal wound-repair signatures (Figure S3E; Tables S1A-G and S2). These findings support the possibility that low-transcript populations reflect dedifferentiated or reserve stem-like states. Overall, mock-treated colonoids exhibited ongoing restitution within a broader inflammation-associated transcriptional landscape.

Finally, we examined whether inflammatory markers associated with CDI localized to specific epithelial populations within mock colonoids. *CXCL8*, *TNF*, *IL1B*, *IL15*, and *CCL20* were distinctly enriched within M-like cells across timepoints (Figure 3F).

### M-like cells define a *CFTR*-linked CDI vulnerability state

Inducible Microfold (M)-like cells arise in the colon during regeneration following epithelial damage and inflammation and have been associated with *CCL20*-dependent differentiation.^56^ Therefore, we asked whether the M-like population in our single-cell dataset shared transcriptional features with other inflammation or regenerative colon epithelial cell types. Relative to all other cell states, the M-like cluster across both Mock and CDI conditions demonstrated strong transcriptional overlap with cell type signatures associated with inflammation, epithelial damage, and repair. These included INFLARE,^69^ LND,^70^ CARSC-like, ^47^ SPIB⁺ inflamed epithelial,^58^ IAF,^56^ and Paneth-like cell types^71^ (Figure 4A and Figure S4A).

**Figure 4.**
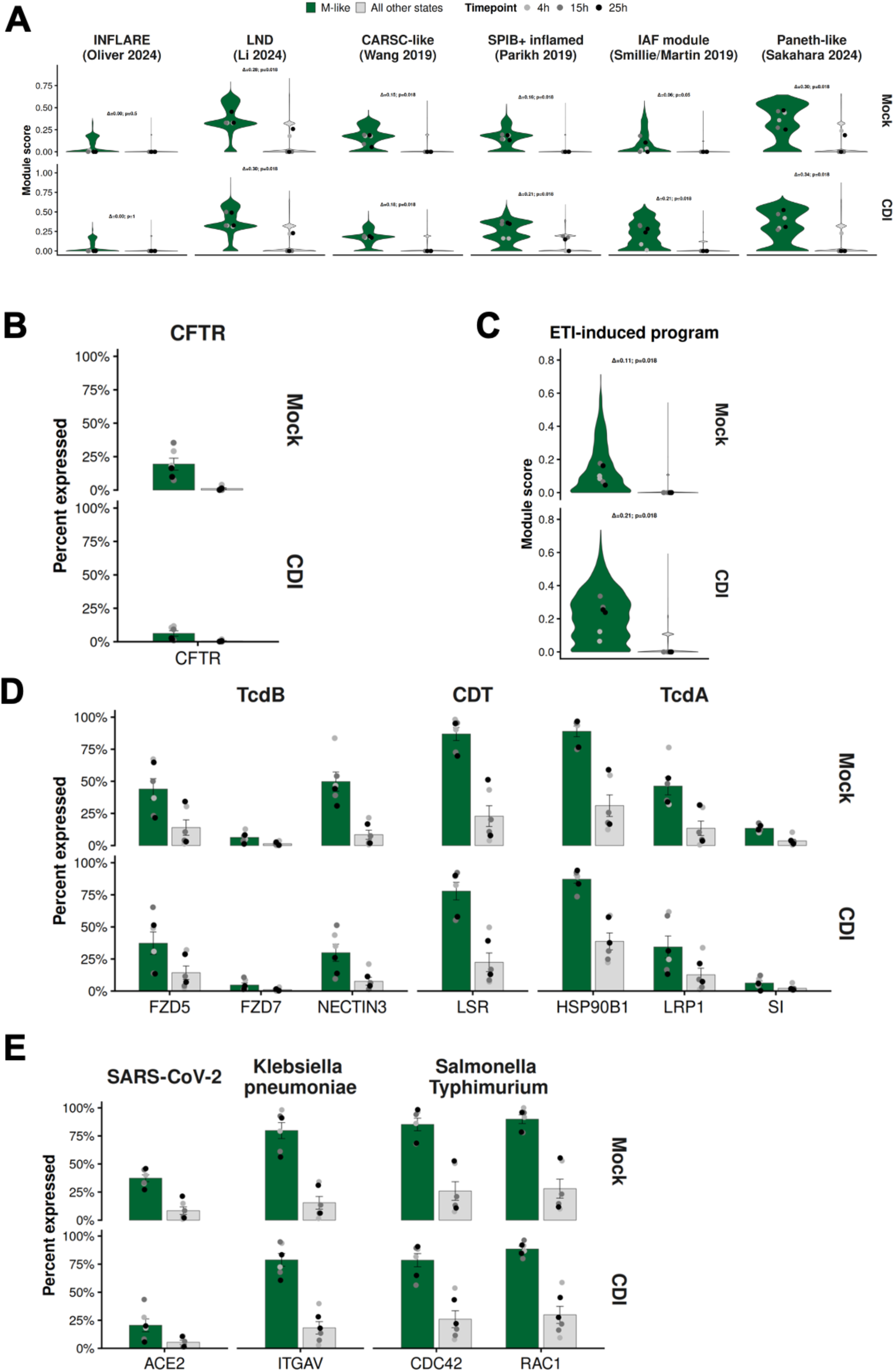
M-like cells define a *CFTR*-linked CDI vulnerability state. (A) Violin plots of module scores for INFLARE, LND, CARSC-like, SPIB⁺ inflamed epithelium, IAF, and Paneth-like transcriptional programs shown separately for Mock and CDI conditions. Green denotes M-like cells and gray denotes all other cell states combined. Points represent replicate–timepoint medians (4 h, 15 h, 25 h). Δ indicates the mean replicate-level difference between M-like and all other states within Mock and CDI samples; p values were calculated using one-sided Wilcoxon signed-rank tests. (B) Percent *CFTR* expressing cells shown separately for Mock and CDI samples. Bars represent mean percent of expressing cells across replicate–timepoint units and error bars indicate standard error. Points represent individual replicate–timepoint values. (C) Violin plots of ETI-induced transcriptional program scores shown separately for Mock and CDI samples. Plot formatting and statistical annotations are displayed as in panel A. (D) Percent expressing cells of *C. difficile* toxin receptor and host-interaction genes grouped by toxin class. TcdB-associated genes include *FZD5*, *FZD7*, and *NECTIN3*; CDT receptor genes include *LSR*; and TcdA-associated factors include *HSP90B1*, *LRP1*, and *SI*. Bars and points are summarized as in panel B. (E) Percent expressing cells of additional epithelial host-interaction genes for other pathogens, including *ACE2* (SARS-CoV-2), *ITGAV* (*Klebsiella pneumoniae*), and *CDC42*/*RAC1* (*Salmonella Typhimurium*). Bars and points are summarized as in panel B.

**Figure S4.**
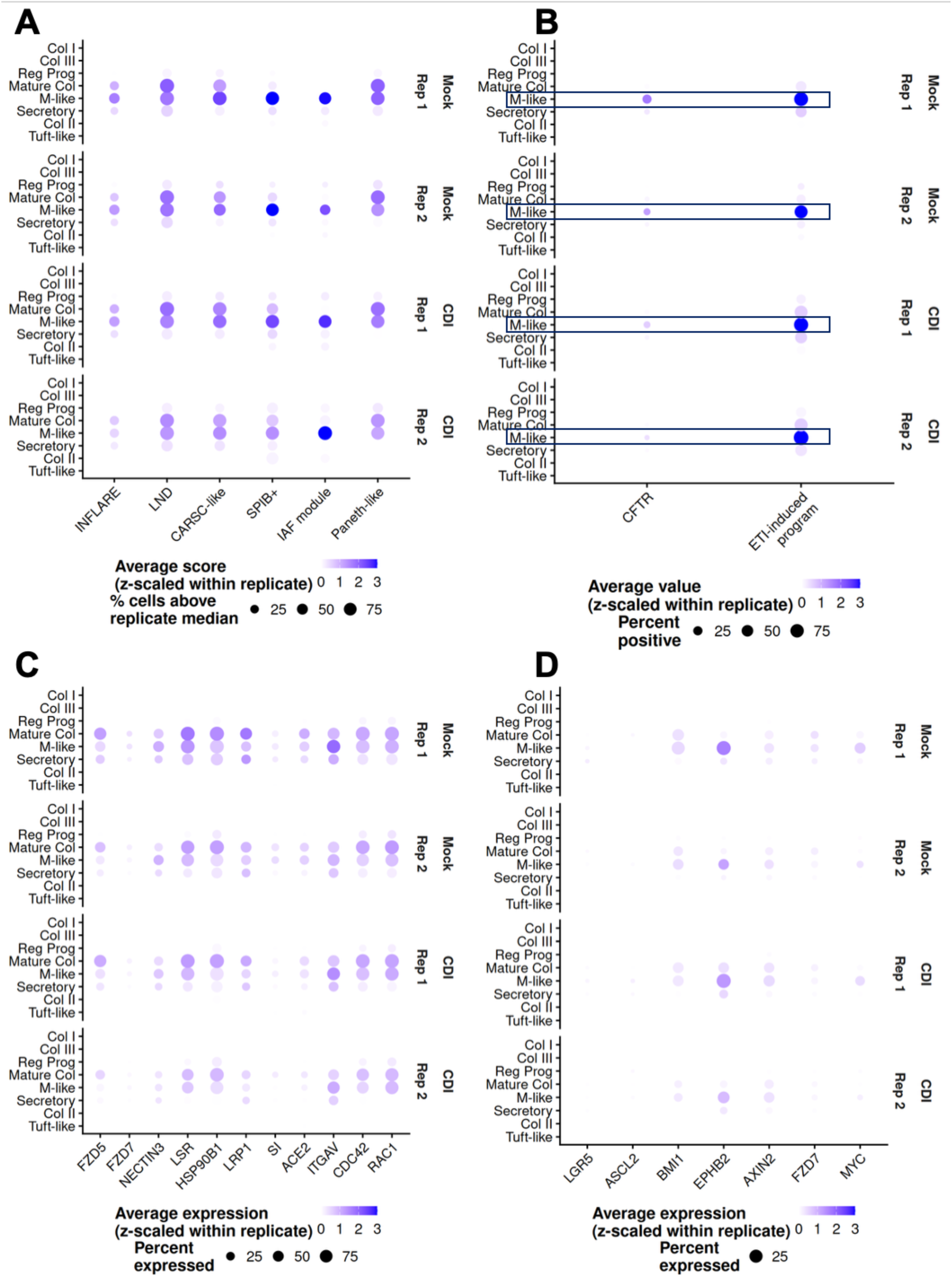
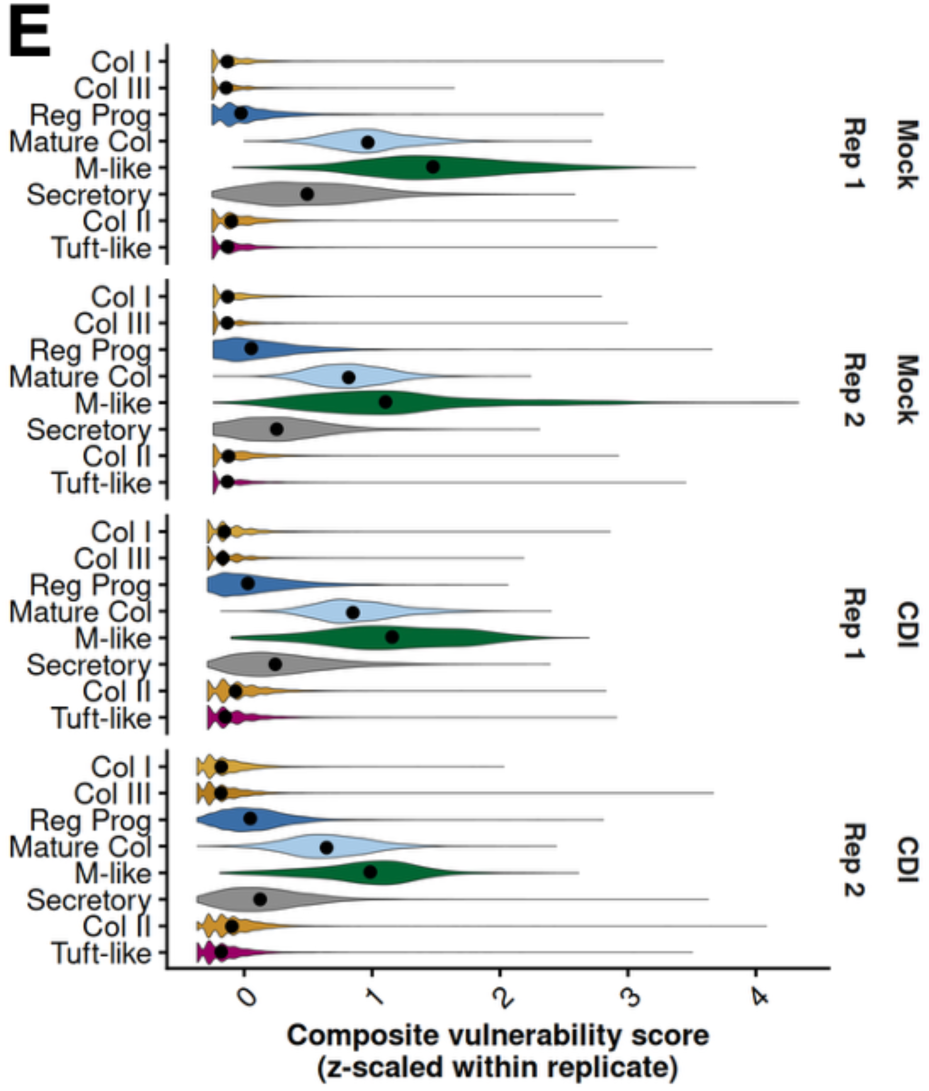
Regenerative, CFTR-associated, and host-interaction programs across epithelial cell states in Mock and CDI colonoid replicates, related to Figure 4. (A) Dot plot of regenerative and inflammation-associated transcriptional programs across epithelial cell states in Mock and CDI replicate samples. Dot size represents the percentage of cells above the replicate-specific feature median and color indicates average score (z-scaled within replicate). (B) Dot plot of CFTR expression and ETI-induced transcriptional program scores across epithelial cell states in Mock and CDI replicate samples. Dot size represents the percentage of positive cells within each biological replicate and color indicates average expression (z-scaled within replicate). (C) Dot plot of host-interaction genes across epithelial cell states in Mock and CDI conditions. Genes include C. difficile toxin receptors and host-interaction factors (FZD5, FZD7, NECTIN3, LSR, HSP90B1, LRP1, SI) together with epithelial interaction genes associated with other pathogens (ACE2, ITGAV, CDC42, RAC1). Dot size represents the percentage of positive cells within each biological replicate and color indicates average expression (z-scaled within replicate). (D) Dot plot of stem-cell and Wnt signaling markers across epithelial cell states in Mock and CDI replicate samples. Markers were selected from Mileto et al. study of CDI-associated stem-cell damage and epithelial repair dysfunction. Dot size represents the percentage of positive cells within each biological replicate and color indicates average expression (z-scaled within replicate). (E) Violin plots of composite vulnerability scores across epithelial cell states shown separately for Mock and CDI samples. Scores were normalized independently within biological replicates before visualization. Points represent replicate–timepoint medians (4 h, 15 h, 25 h).

This finding is notable in the context of cystic fibrosis, which is characterized by chronic inflammation and impaired CFTR-dependent regenerative capacity. Therefore, we examined *CFTR* expression across colonoid clusters. *CFTR*-expressing cells were almost completely confined to the M-like cluster in both Mock and CDI conditions (Figure 4B and Figure S4B). For further evaluation of a possible link between the M-like cells and *CFTR*-associated biology, we derived a transcriptional program generated from published intestinal organoid datasets of patients with cystic fibrosis before and after pharmacologic *CFTR* correction with elexacaftor/tezacaftor/ivacaftor (ETI). This ETI-treatment profile was strongly enriched within the M-like cluster under both Mock and CDI conditions, with lower enrichment observed in epithelial states along the M-like lineage trajectory (Figure 4C and Figure S4B). This links *CFTR*-associated transcriptional signatures to the epithelial state most strongly enriched for CDI vulnerability markers.

Because CDI susceptibility depends on epithelial host–pathogen interactions, we mapped genes implicated in *C. difficile* toxin and target interaction. Several toxin receptors and host-interaction factors, including *FZD5*, *FZD7*, *NECTIN3*, *LSR*, *HSP90B1*, *LRP1*, and *SI*, were detected within M-like lineage clusters across CDI status (Figure 4D and Figure S4C). Similar expression patterns were observed for genes linked to other pathogen infections associated with colonic inflammation, including *ACE2* (SARS-CoV-2), *ITGAV* (*Klebsiella pneumoniae*), and *CDC42* and *RAC1* (*Salmonella Typhimurium*). Enrichment was consistently highest within the M-like lineage (Figure 4E and Figure S4C), further linking this cell state to host–pathogen interaction programs.

Prior work has shown that severe CDI can disrupt epithelial repair through toxin-mediated injury to the colonic stem-cell compartment.^16^ In line with these observations, M-like cells in our model were enriched for markers previously linked to CDI-induced stem-cell dysfunction, including *BMI1*, *EPHB2*, *AXIN2*, and *MYC* (Figure S4D).

To integrate these findings, we computed a composite vulnerability score incorporating regenerative, inflammatory, CFTR-associated, and host-interaction programs. In Mock colonoids, positive scores were restricted to the M-like lineage and peaked within the M-like state overall (Figure S4E). In CDI, positive vulnerability scores also emerged within the tuft-like lineage (Figure S4E), suggesting broader epithelial involvement during infection response and repair.

### CDI reshapes epithelial repair programs through a shift from M-like to tuft-like lineage involvement

We next examined how CDI altered transcription across epithelial lineages. M-like cells increased in number within the M-like cluster during infection, particularly at 25 h post inoculation (Figure 5A). Under Mock conditions, progression along the M-like lineage was associated with a highly transcriptionally active M-like population (Figure S5A). In response to CDI, however, the M-like cluster and neighboring lineage populations became increasingly enriched for lower-transcript Hectoscript cells (Figure S5A). This shift coincided with reduced representation of cells with high mitochondrial read ratios (≥40%) (Figure S5B-C). In Mock samples, high mito-ratio cells within the M-like lineage were composed primarily of Kiloscript cells, whereas CDI-associated high mito-ratio populations became enriched for Hectoscript cells before disappearing at later timepoints (Figure S5C). Consistent with disrupted progression through this regenerative trajectory, CDI-associated M-like cells also showed reduced *CFTR* expression, and a lower proportion of *CFTR*-expressing cells compared with Mock cell profiles (Figures 4B and S4B).

**Figure 5.**
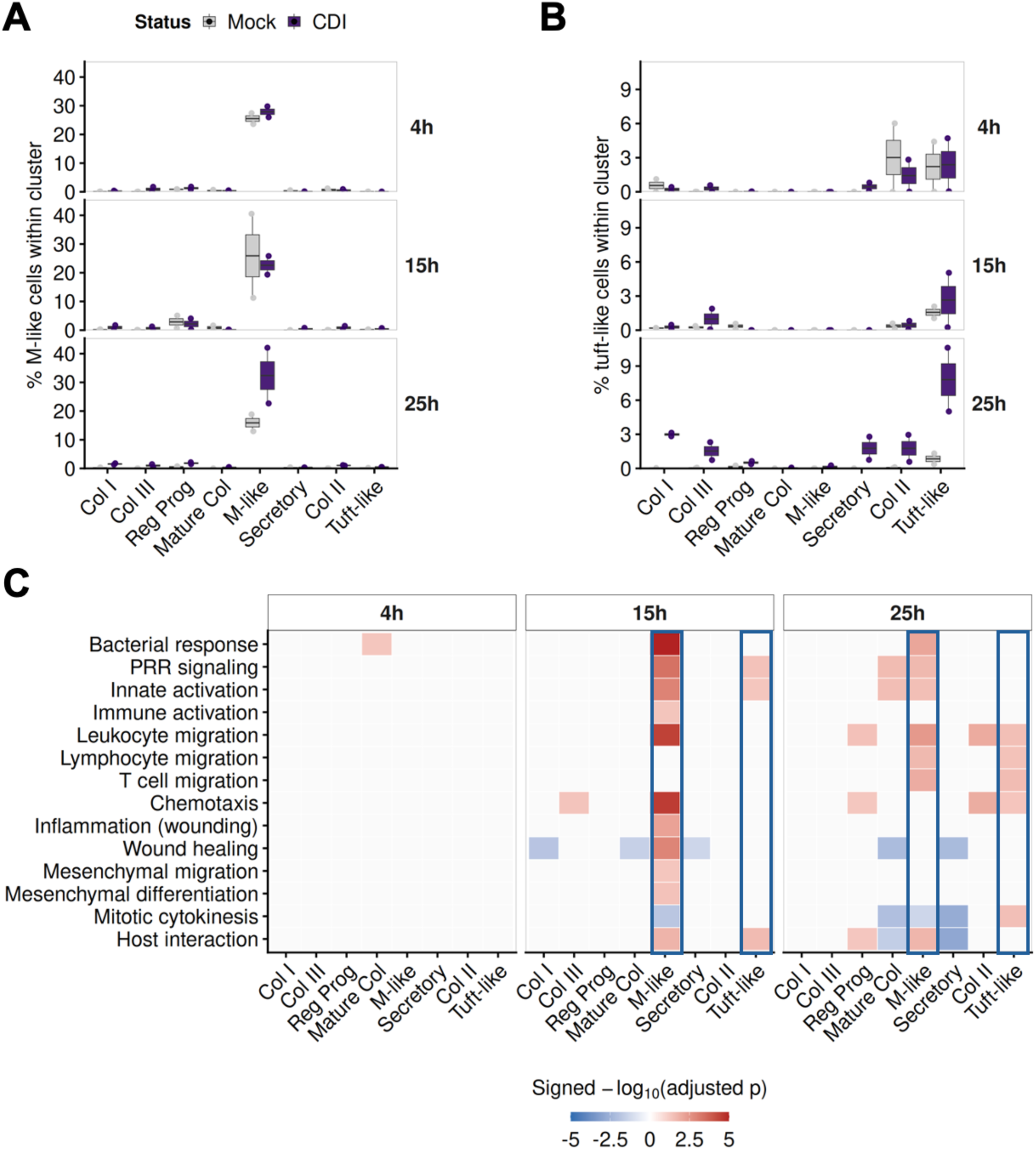
CDI reshapes epithelial repair programs through a shift from M-like to tuft-like state involvement. (A) Percentage of M-like cells within each cell state cluster across timepoints in Mock and CDI samples. (B) Percentage of tuft-like cells within each cell state cluster across timepoints in Mock and CDI samples. (C) Heatmap of selected *CCL20*-associated GO across cell states and timepoints. Values represent −log10(adjusted p), with red indicating enrichment among genes upregulated in CDI and blue indicating enrichment among genes downregulated in CDI.

**Figure S5.**
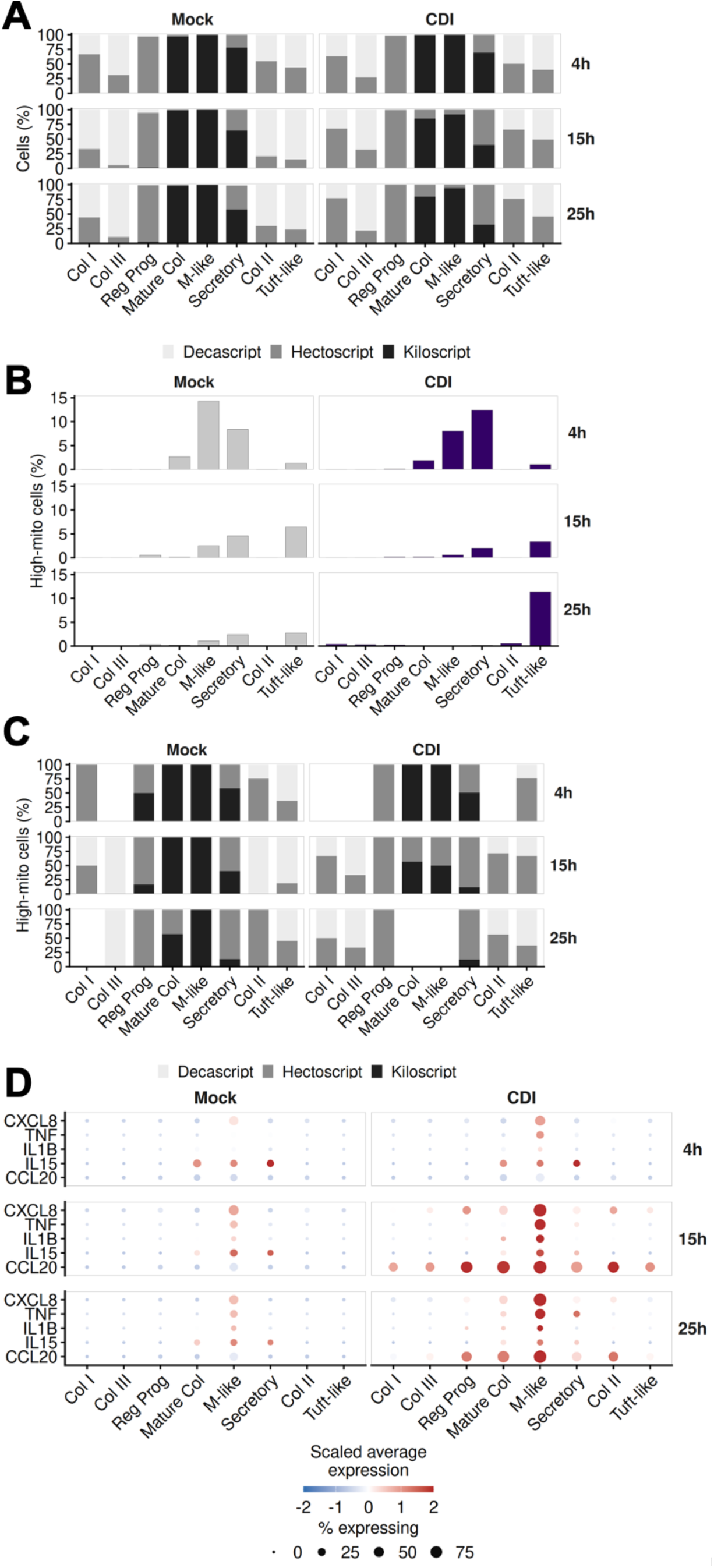
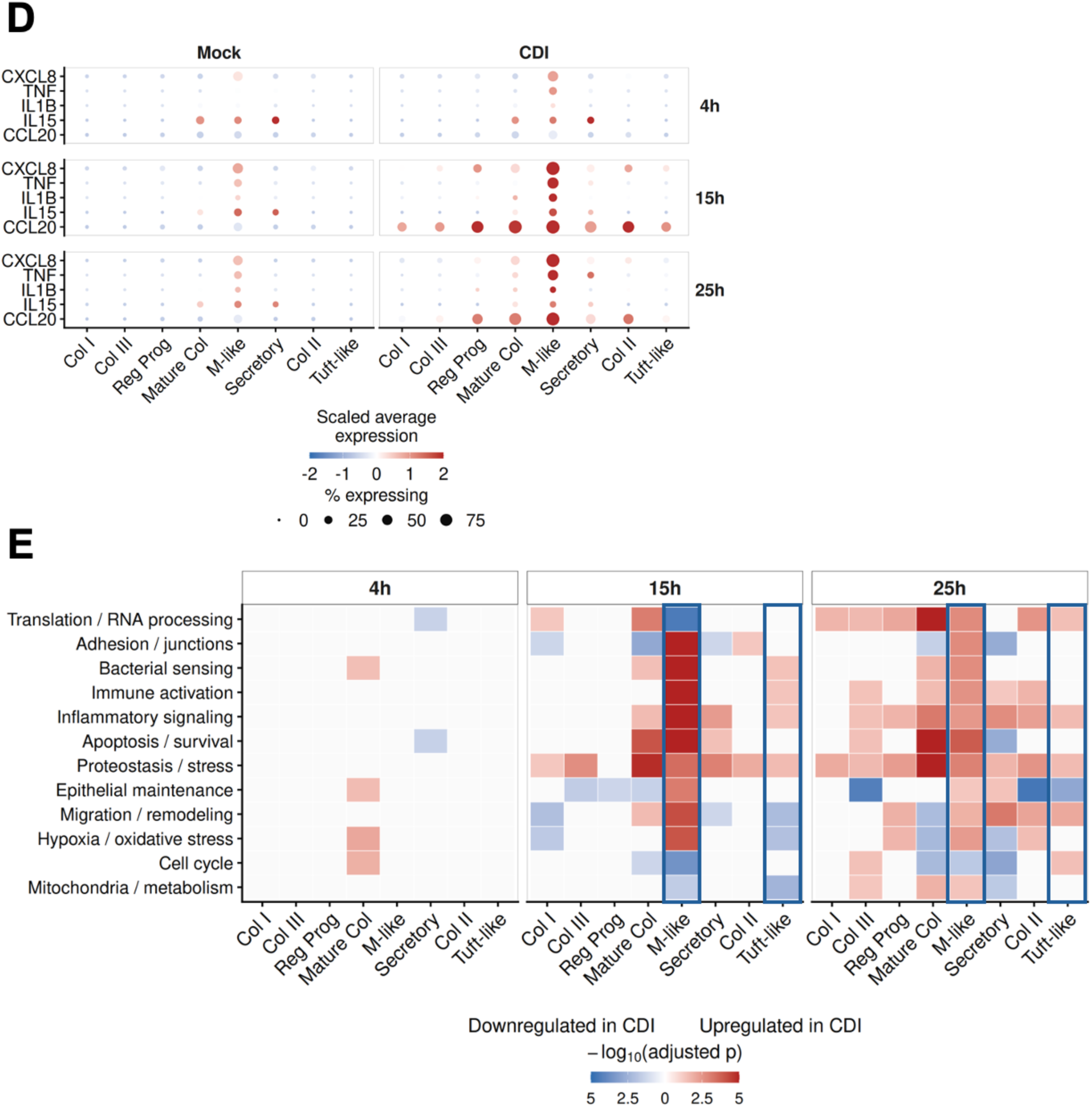
Responses by cell state to CDI in transcript complexity, mitochondrial burden, and injury-response programs, related to Figure 5. (A) Distribution of Decascript, Hectoscript, and Kiloscript cells across cell states, timepoints, and samples. (B) Percentage of high-mito cells, defined as cells with mitochondrial read ratio >40%, across cell states, timepoints, and samples. (C) Distribution of Decascript, Hectoscript, and Kiloscript classes among high-mito cells across cell states, timepoints, and samples. (D) Dot plot of *CXCL8*, *TNF*, *IL1B*, *IL15*, and *CCL20* expression across cell states, timepoints, and samples. Dot size indicates the percentage of cells expressing each gene and color indicates scaled average expression. (E) Heatmap of representative GO categories after excluding GO associated with *CCL20*. For each category, cell state, and timepoint, the most statistically significant GO term is shown. Values represent−log10(adjusted p), with red indicating enrichment among genes upregulated in CDI and blue indicating enrichment among genes downregulated in CDI.

By contrast, tuft-like cells expanded broadly across the tuft-like lineage in response to infection (Figure 5B). Under Mock conditions, tuft-like lineage populations were composed predominantly of Decascript cells, and this fraction increased over time (Figure S5A). CDI-associated tuft-like populations did not show this increase. Tuft-like lineage populations also became increasingly enriched for high mito-ratio transcriptomes over time in both Mock and CDI conditions, although the associated transcript-count compositions differed substantially (Figure S5B-C and Figure S1B). These populations overlapped with regions of elevated CytoTRACE2 scores (Figure S3D), consistent with increased plasticity. At later timepoints, tuft-like lineage populations also showed increased expression of *CCL20* (Figure S5D).

Because *CCL20* was among the most upregulated genes induced by CDI, we examined selected *CCL20*-associated GO enrichment across clusters and timepoints (Figure 5C, Tables S3–S4). At 15 hpi, bacterial response and innate immunity programs were concentrated within the M-like lineage, while neighboring lineage populations showed reduced enrichment of wound repair programs (Figure 5C). By 25 h, genes associated with mitotic cytokinesis were downregulated within the M-like lineage but upregulated in the tuft-like cluster (Figure 5C).

Beyond *CCL20*-associated programs, gene expression changes emerged as early as 4 h post inoculation (Figure S5E). Within the colonocyte cluster immediately preceding the M-like population along the lineage trajectory, there was increased enrichment of cell cycle, bacterial sensing, and stress response programs. Clusters immediately following the M-like population along the inferred trajectory showed downregulation of translation and survival-associated pathways.

### CDI amplifies epithelial damage and CCL20 signaling at wound-like sites

Because CCL20 localizes to sites of epithelial injury and repair, we examined the spatial organization of CCL20+ epithelial cells in colonoid monolayers. Immunofluorescence imaging revealed CCL20-producing cells surrounding wound-like regions within the Mock and CDI monolayers. In Mock conditions, CCL20 signal was relatively low and localized to small, discrete epithelial gaps (Figure 6A). In contrast, CDI monolayers exhibited a significantly greater number of wound-like gaps (Figure 6B). The CCL20 signal remained associated with these regions but was markedly increased in intensity and extended into the surrounding areas compared with Mock conditions (Figures 6A and 6C).

**Figure 6.**
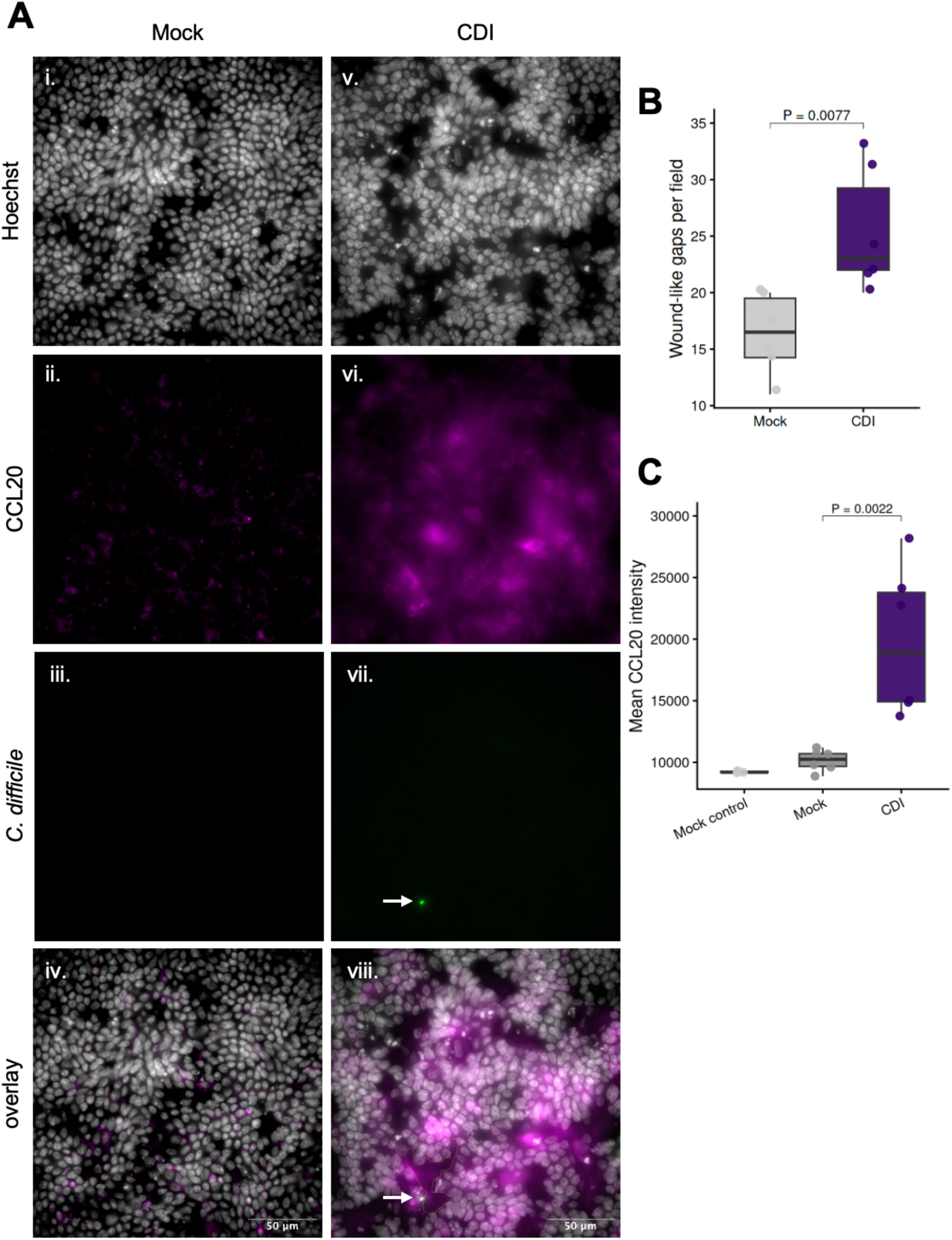
CDI amplifies epithelial damage and CCL20 signaling at wound-like sites. (A) Representative immunofluorescence images of colonoids under Mock and CDI conditions. Nuclei are labeled with Hoechst (i, v), CCL20 is shown in magenta (ii, vi), and *C. difficile* is shown in green (iii, vii). Overlay images are shown in (iv, viii). Scale bars, 50 μm. Arrows indicate representative *C. difficile* bacterium near epithelial surface. (B) Quantification of wound-like gaps per imaging field in Mock and CDI samples. Each point represents an individual imaging field. Boxes indicate median and interquartile range. P value was calculated using a Wilcoxon rank-sum test. (C) Mean CCL20 intensity measured across mock control regions, regions adjacent to mock wound-like gaps and CDI wound-like gaps. Each point represents an individual imaging field. Boxes indicate median and interquartile range. P value was calculated using a Wilcoxon rank-sum test.

Detection of *C. difficile* near the epithelial surface was uncommon in the imaging fields examined. However, when bacteria were observed, they appeared adjacent to disrupted epithelial regions (Figure 6A). Together, these findings indicate that CDI is associated with persistent epithelial disruption accompanied by increased CCL20 signaling at wound-like regions.

## DISCUSSION

Our results demonstrate that *C. difficile* infection does not impose a new transcriptional program in epithelial cells but instead perturbs active pathways of injury and regeneration. We observed inflammatory and regenerative epithelial populations in colonoid monolayers with and without infection. This informed our analytical strategy, including use of disease-specific epithelial references for cell-type annotation and retention of low-transcript epithelial populations commonly excluded during conventional scRNA-seq filtering, enabling reconstruction of regenerative trajectories. By mapping the single-cell landscape of human colonoids, we identified an inducible M-like cell type that emerges during colonic inflammation and epithelial repair and is enriched for CDI-linked inflammatory, CFTR, and toxin-interaction programs within a single epithelial population. Early infection responses localized to the M-like lineage, followed by expansion of tuft-like populations associated with increased plasticity, altered transcript complexity, and mitochondrial remodeling. Together, these findings suggest that epithelial regeneration creates a permissive environment for CDI while infection disrupts restoration of epithelial integrity.

Canonical M cells, established entry points for various intestinal pathogens, are constitutively present in the healthy small intestine, whereas colonic M-like populations emerge and expand with increasing epithelial inflammation and repair.^72–75^ Because *C. difficile* exploits the inflamed colon to promote virulence, the damaged epithelium may provide a niche in which vulnerable epithelial populations and permissive conditions coexist.^76^ This framework may help explain the increased susceptibility to CDI and severe disease in patients with high levels of pre-infection colonic inflammation, including IBD and antibiotic-treated individuals.

Colonic inflammation alone, however, is insufficient to dictate CDI susceptibility. Patients with cystic fibrosis (CF) experience chronic inflammation, frequent antibiotic exposure, and high rates of *C. difficile* colonization, yet symptomatic CDI remains uncommon. CF is characterized by defective CFTR signaling and impaired regeneration, whereas treatment with the CFTR corrector ETI accelerates epithelial wound repair.^77^ In our colonoid system, CFTR expression was largely restricted to the M-like cluster, where the ETI-associated transcriptional signature was also most enriched. During CDI, however, CFTR expression within this population was reduced, raising the possibility that M-like cells and CFTR-associated repair programs are targeted during infection. A published case report describing symptomatic CDI following ETI initiation in a patient with CF previously exposed to antibiotics without CDI further aligns with this possibility.^78^

Similarly, the relative protection from symptomatic CDI observed in neonates may reflect impaired regenerative capacity. Neonatal protection has historically been attributed to lack of toxin receptor expression. However, our results show that toxin receptors and target genes are concentrated within regenerative M-like epithelial populations. Notably, newborns do not produce mature M cells in the small intestine, potentially limiting formation of this CDI vulnerable epithelial lineage.^79,80^

Prior work has shown that CDI is more severe when toxins gain access to the stem cell compartment causing damage and disrupting epithelial repair.^16^ Here, we refine this model by suggesting that CDI vulnerability emerges during regeneration itself, without infection. In our colonoid monolayers, wound-like gaps surrounded by CCL20-producing epithelial cells were present even in the absence of infection. Since *CCL20* is a marker for M cells in the inflamed IBD colon and increases at sites of epithelial repair, our baseline culture conditions appeared to be actively undergoing restitution.^56,81,82^ In the scRNA-seq dataset, M-like cells demonstrated the highest and most selective expression of *CCL20* among epithelial populations, suggesting that the CCL20+ cells localized near wound-like regions may correspond to this regenerative cell type. During CDI, CCL20 production became broader and more intense while remaining concentrated near sites of epithelial disruption. Together with transcriptional evidence of downregulated wound repair programs preceding the M-like population, it appears that CDI amplifies pre-existing regenerative signaling while preventing successful epithelial repair. This disruption may be driven in part by toxin activity, as toxin-deficient strains induce substantially less damage in vivo, and our prior work demonstrated that toxin activity is required for robust epithelial CCL20 induction.^64^

Markers of epithelial populations previously found to be targeted by *C. difficile* toxins were most highly expressed in M-like cells.^16^ Without infection, the M-like lineage was associated with epithelial restitution, whereas CDI shifted epithelial responses toward tuft-like populations, possibly as a compensatory response. Prior studies showing tuft cell depletion by days 3–5 post-infection suggest that these cells may later become secondary targets during repair.^83^ Recurrent CDI may therefore reflect cyclical disruption of epithelial restitution in which initial infection depletes vulnerable M-like populations, followed by tuft-like expansion and subsequent targeting during compensatory repair. During recovery, regeneration may transiently restore M-like populations, recreating permissive epithelial states associated with recurrent disease 2–4 weeks after treatment.^47^ Repeated cycles of injury and incomplete repair could progressively impair epithelial restitution, leading to chronic inflammation and abnormal persistence or accumulation of specialized epithelial populations, including M-like and tuft-like cells, as observed in IBD.

Although baseline injury and inflammation were not intentionally induced in this study, mock-treated colonoids exhibited epithelial damage and antibiotic response programs that may relate in part to antibiotic exposure in cell culture. Primocin alters stem cell transcriptional programs and mitochondrial metabolism, and shares mechanistic similarities with clindamycin, a high-risk CDI-associated antibiotic.^84^ Under these conditions, colonoids may model aspects of the inflamed and regenerative epithelium associated with increased CDI susceptibility.

Finally, the CDI vulnerability identified in the M-like lineage may extend beyond CDI. This population preferentially expressed host factors linked to interactions with other pathogens, including SARS-CoV-2, *Salmonella Typhimurium*, and *Klebsiella pneumoniae*. Thus, epithelial regenerative state may be a more precise determinant of infection vulnerability and disease severity than colonic inflammation alone.

## STUDY LIMITATIONS

This study has several limitations. The colonoid monolayer model consists of epithelial cells only and does not incorporate immune or other cell types, or microbiota that shape host responses in vivo. Our conclusions are based primarily on transcriptional associations, rather than direct functional analyses of specific epithelial states. In addition, regenerative trajectories inferred from snapshot single-cell transcriptomes represent modeled transitions rather than direct observation of cell-state progression over time. Finally, antibiotic-containing colonoid culture conditions are not equivalent to clinical antibiotic exposure, and parallels between these systems should be interpreted cautiously.

## DATA AVAILABILITY

The raw and processed scRNA-seq data from this study have been deposited to the NCBI Gene Expression Omnibus (GEO) under accession number GSE331313 and will be made available upon publication.

## METHODS

### Preparation and Differentiation of Colonoid Monolayers

Human colonoids, derived from the colonoid cell line CJ50, were cultured and differentiated on transwell monolayers as previously described.^64^ Colonoids were grown in Matrigel using 65% L-WRN medium, with Primocin and 5 μM Y-27632 added during the first 48 hours of culture.

Dissociated colonoids were seeded onto Matrigel-coated 6.5 mm Transwell inserts (0.4 μm pore size; Costar 3413) and expanded to confluence. Confluent monolayers were differentiated for 5 days in medium containing Primocin prior to inoculation with pre-germinated *C. difficile* spores. Transepithelial electrical resistance (TEER) measurements were obtained using a Millicell ERS-2 volt/ohm meter according to the manufacturer’s instructions.

### Clostridioides difficile inoculation

Pregerminated *C. difficile* UK1 spores were added to the apical compartment of differentiated colonoid monolayers as previously described.^64^ Mock-treated monolayers received buffer only. Monolayer cells were harvested at 4, 15, and 25 h post inoculation.

### Single cell RNA sequencing

For single-cell dissociation, monolayers were washed with PBS containing 0.5 mM EDTA and incubated with TrypLE Express at 37°C. Dissociated cells were filtered through a 40 μm cell strainer, resuspended in 0.04% BSA containing Y27632, and counted prior to library preparation.

Single-cell libraries were generated using the Chromium Next GEM Single Cell 3′ v3.1 platform (10x Genomics) according to the manufacturer’s instructions. Library quality was assessed using an Agilent Bioanalyzer, and paired-end sequencing was performed by the Tufts University Genomics Core Facility.

## COMPUTATIONAL METHODS

### scRNA-seq analysis

Raw sequencing data were pre-processed using Cell Ranger (10x Genomics). Seurat v5.0.2 was used for scRNA-seq analyses.^85^ Cells with >75 detected genes were retained. Mock and CDI samples for all timepoints and replicates were normalized independently using SCTransform and integrated using Harmony integration method. Principal component analysis was performed. Clustering was performed using the Louvain algorithm at resolution 0.4.

### Cell annotation and cluster labeling

Single-cell profiles were annotated using SingleR with the Smillie et al. human colon single cell atlas as a reference. Initial annotation used combined healthy, non-inflamed IBD, and inflamed IBD colon epithelial datasets. Final annotations were generated using the inflamed IBD epithelial subset. Cell type composition and transcript count distribution were used to label clusters.

### External dataset analysis

Publicly available human colon epithelial single cell RNA-seq datasets from Triana et al.^20^ and Parikh et al.^58^ were analyzed using the same SingleR annotation framework applied to CDI colonoids.

### Crypt-axis and CytoTRACE2 analysis

Crypt-axis scores were calculated using UCell module scoring with a predefined crypt-top epithelial gene set consisting of *SELENOP*, *CEACAM7*, *PLAC8*, *CEACAM1*, *TSPAN1*, *CEACAM5*, *CEACAM6*, *IFI27*, *DHRS9*, *KRT20*, *PKIB*, *HPGD*, *LYPD8*, and *RHOC*.^58^

Developmental potential was estimated using CytoTRACE2.^86^ To account for replicate structure, the integrated Seurat object was split by replicate, and CytoTRACE2 was run separately on each replicate using the human model with default settings except for batch size parameters set to 10,000 cells for prediction and 1,000 cells for smoothing. CytoTRACE2 scores were then retained in the Seurat metadata, replicate objects were merged, and the combined dataset was re-normalized, re-integrated with Harmony, reclustered, and visualized by UMAP for downstream analyses.

### RNA velocity

RNA velocity analysis was performed on mock-treated epithelial cells using velocyto-generated spliced and unspliced transcript matrices derived from Cell Ranger alignment files. Filtered cell barcodes from the Seurat object were matched to velocyto loom files for each biological replicate, and spliced and unspliced matrices were incorporated as separate assays. Velocity estimates were calculated using SeuratWrappers and velocyto.R with PCA-based nearest-neighbor smoothing (kCells = 25, fit.quantile = 0.02) and projected onto Harmony-integrated UMAP embeddings.

### Trajectory inference and pseudotime analysis

Trajectory inference was performed using Slingshot on Harmony-integrated UMAP embeddings.^87^ Cells were grouped using simplified cluster labels, and trajectories were initialized using the Colonocyte I population as the starting cluster. Lineages were inferred using simultaneous principal curves with shrinkage and reweighting enabled (shrink = 0.10, reweight = TRUE, stretch = 2).

Pseudotime values were independently scaled from 0 to 1 within each lineage. Cells lacking assigned pseudotime values for a given lineage were excluded from lineage-specific analyses. Cell fraction in each cluster across pseudotime was summarized by binning scaled pseudotime values into equal intervals and calculating the proportion within each bin.

Sensitivity analyses were performed using Colonocyte III and regenerative progenitor populations as alternative starting clusters.

### Module scoring

Published epithelial injury, inflammation, and repair-associated gene programs were evaluated using UCell module scoring. Gene sets included INFLARE, LND, inflammatory fibroblast-associated, Paneth-like, and ETI/CFTR-associated signatures. Only genes present in the final Seurat object were included. The INFLARE signature included *MUC5AC*, *AQP5*, and *BPIFB1*. The LND signature included *LCN2*, *NOS2*, and *DUOX2*. The inflammatory fibroblast-associated signature included *IL6ST*, *IL11*, *CXCL2*, *CXCL8*, *CXCL1*, *CXCL5*, *CCL2*, and *CCL7*. The Paneth-like signature included *LYZ*, *NOD2*, and *CD24*. The ETI/CFTR-associated signature included *AMOTL2*, *ANKRD1*, *BNIP5*, *CCN1*, *CCN2*, *CXCL3*, *CXCL8*, *DUSP5*, and *EDN1*. A CDI inflammatory score was calculated using *CXCL8*, *TNF*, *IL1B*, and *IL15*.

### Differential expression and Gene Ontology analysis

Differential expression analysis was performed in Seurat using Wilcoxon rank-sum testing (FindMarkers). Comparisons between CDI and mock-infected single-cell profiles were performed independently within cluster, matched timepoint and biological replicate. Genes detected in at least 5% of cells were included (min.pct = 0.05), and testing used an initial log fold-change threshold of 0.10. Significant differentially expressed genes were defined as adjusted P < 0.05 and absolute log2 fold-change ≥ 0.25. Replicate concordant differentially expressed genes were identified by intersecting significant genes detected independently in both biological replicates while requiring concordant direction of expression change.

Gene Ontology enrichment analysis was performed using clusterProfiler with Biological Process ontology classification. Gene symbols were converted to Entrez identifiers using org.Hs.eg.db. Enriched GO terms were filtered using Benjamini–Hochberg-adjusted thresholds (p.adjust < 0.05, qvalue < 0.20). Related GO terms were grouped into broader biological categories for visualization.

### Immunofluorescence staining and imaging

Colonoid monolayers were fixed in 4% paraformaldehyde, permeabilized with Triton X-100, and blocked prior to staining as previously described.^64^ Monolayers were stained using antibodies against CCL20 and *C. difficile* together with Hoechst nuclear stain. Images were acquired using a Leica DMi8 fluorescence microscope and processed in Fiji/ImageJ.

### Quantification of wound-like gaps and CCL20 intensity

Immunofluorescence images were analyzed in Fiji/ImageJ. A restricted Z range spanning the epithelial layer was selected using the nuclear channel and projected using the same projection method across channels. Nuclei-depleted epithelial regions were defined on the nuclear channel as contiguous regions within the epithelial sheet lacking nuclear staining and exceeding normal intercellular spacing. Regions at image borders, out-of-focus areas, imaging artifacts, and small spaces between individual cells were excluded.

Regions of interest were drawn manually on the nuclear channel and transferred to matched CCL20 images for quantification. ROI area, mean CCL20 fluorescence intensity, and integrated density were measured in Fiji/ImageJ. Mean fluorescence intensity was used for comparisons.

For field-level analyses, CCL20 intensity surrounding nuclei-depleted regions was measured and averaged within each field and compared with averaged intact epithelial control regions from the same field. Local CCL20 enrichment was calculated as the ratio of mean CCL20 intensity in nuclei-depleted regions relative to intact epithelial regions. The number of nuclei-depleted regions per field was also quantified.

Visible *C. difficile* signal was assessed qualitatively within the matched bacterial channel to determine whether bacterial signal localized adjacent to nuclei-depleted epithelial regions.

### Statistical analysis

Statistical analyses were performed in R. Wilcoxon rank-sum testing and Benjamini–Hochberg multiple testing correction were used where indicated. Replicate-level summaries were used for comparative analyses and visualization.

## ACKNOWLEDGEMENTS

The authors thank Drs. Aimee Shen, Joan Mecsas, Bree Aldridge, Sergio Triana, and Alex Shalek for generously providing helpful suggestions and expertise. The authors acknowledge the Tufts University High Performance Compute Cluster, which was utilized for some of the research reported in this paper.

